# Antitrypsin surrogate, Alphataxin, increases tumor CD4^+^ T cells and suppresses murine colon cancer

**DOI:** 10.64898/2026.04.06.716656

**Authors:** Cynthia L. Bristow, Thomas Q. Garvey, Ronald Winston

## Abstract

CD4^+^ T helper cells are required for CD8^+^ killer T cells to suppress tumor growth. An orally-available small molecule surrogate of alpha-1 antitrypsin, Alphataxin, was previously demonstrated to elevate the numbers of circulating and tumor-infiltrating CD4^+^ T cells and to suppress kidney tumor growth in mice. To determine whether Alphataxin might be effective in other T cell-responsive cancers, mice orthotopically implanted with colon tumors were treated using Alphataxin and anti-PD-1 as monotherapies or in combination. Combination therapy significantly suppressed tumor growth (ORR = 37.5%) and increased tumor-infiltrating CD4^+^ T cells, CD8^+^ T cells, NK cells, M2 macrophages, and DC2 dendritic cells. Release of IFN-γ by helper T cells in the tumor microenvironment appeared to contribute to the effectiveness of killer T cells in suppressing tumor growth. Toxicology studies in rats revealed no untoward effects. Alphataxin, to our knowledge the first and only drug developed to rapidly and sustainably increase the number of circulating and tumor-infiltrating CD4^+^ helper T cells, is a powerful therapeutic that provides long-term remission in T cell-responsive cancers in combination with anti-PD-1.

## INTRODUCTION

In two small clinical trials, we previously showed that treatment with the FDA-approved plasma protein alpha-1 antitrypsin (AAT, alpha-1 proteinase inhibitor) caused a dramatic, sustained increase in circulating numbers of functionally competent CD4^+^ T cells and/or increased CD4/CD8 T cell ratios (Bristow *et al*, 2010; Bristow *et al*, 2019). In contrast, numbers of CD8^+^ T cells, monocytes, or granulocytes were not increased (Bristow *et al*., 2010). Binding of AAT to plasma membrane-associated leukocyte elastase induces aggregation of functionally-related receptors that include CD4, TcR, chemokine receptors, and their cargo which subsequently bind to members of the low density lipoprotein receptor family causing endocytosis and NFκB signaling (Bristow *et al*, 2003; Bristow *et al*, 2013). This activity occurs as an epiphenomenon of cellular locomotion through tissue including thymus from which immature T cells emerge as mature T cells (Bristow *et al*., 2013; Bristow *et al*, 2021; Bristow & Winston, 2021a, b). We demonstrated in one clinical trial that AAT is required for immature T cells to become mature CD4^+^ T cells (Bristow *et al*., 2019).

We have developed the orally available small molecule Alphataxin to act as a surrogate for AAT. Alphataxin was previously shown to increase the number of circulating and tumor-infiltrating functional CD4^+^ helper T cells in healthy mice and in a murine kidney cancer model (Bristow *et al*., 2021). Alphataxin, significantly suppressed or regressed tumor growth and metastasis as a monotherapy or in combination with anti-PD-1 in a mouse model of kidney cancer with a 66.7% tumor objective response rate (ORR) (Bristow *et al*., 2021).

CD4^+^ helper T cells are now recognized as central to tumor growth suppression, just as they are in all immune system responses (Farber, 2020; Guo *et al*, 2024; Speiser *et al*, 2023; Sun *et al*, 2023; Venkatesh & Fong, 2025). There are at least 6 subsets of CD4^+^ effector T cells including Th1, Th2, Th9, Th17, Th22, and Th-GMCSF, all of which are derived from naïve Th0 T cells (Zielinski). Th1 T cells are CD4^+^ helper T cells that are unique by primarily releasing a pair of cytokines, IFN-γ and TNFα (Lee *et al*, 2021). While IFN-γ can be released by CD8^+^ T cells, NK cells, and sometimes by NKT and antigen presenting cells, IFN-γ is primarily released by Th1 cells (Lee *et al*., 2021). Binding of IFN-γ and TNFα to receptors on CD8^+^ killer T cells induces receptor-mediated internalization that stimulates signaling via the Jak/Stat pathway and, as a consequence, stimulation of gene expression after which the cytokines are degraded by lysosomotropic-sensitive agents (Celada & Schreiber, 1987; Claudinon *et al*, 2007; Ivashkiv, 2018). In other words, in pathologic situations such as in the tumor microenvironment (TME), depletion of IFN-γ and TNFα provides evidence for the activities of CD4^+^ helper (Th1) and CD8^+^ killer T cells.

Herein, we report that in addition to murine kidney cancer, Alphataxin in combination with anti-PD-1 antibody treatment significantly suppressed or regressed orthotopically implanted colon tumor cell growth as compared to monotherapy or untreated controls. Combination therapy, but not monotherapy, increased the number of CD4^+^ T cells, CD8^+^ T cells, NK cells, M2 macrophages, and DC2 dendritic cells as compared to untreated controls. Further, Alphataxin monotherapy significantly decreased levels of IFN-γ in tumor-associated CD4^+^ T cells while there were no changes in IFN-γ levels in tumor-associated CD8^+^ T cells. These results provide new evidence that Alphataxin increases the number of tumor-infiltrating CD4^+^ T cells and that in combination with anti-PD-1 treatment, Alphataxin provides a potent method to suppress tumor growth.

## RESULTS

### Circulating CD4/CD8 ratio is rapidly increased by Alphataxin

We have previously shown that after 2 weeks of daily Alphataxin treatment, the number of circulating CD4^+^ T cells and tumor-infiltrating CD4/CD8 ratio significantly increased in mice (Bristow *et al*., 2021). To examine the effects of a single Alphataxin dose after 24 hours, 5 male and 5 female rats were administered vehicle or Alphataxin (400 mg/kg) by oral gavage. Between pre-dose and 24 hr post-dose in the vehicle treated control group, there was a significant increase in CD4^+^ T cells (*P* < 0.05), CD8+ T cells (*P* < 0.007), and CD4/CD8 ratio (*P* < 0.0002) (**Fig 1**). These results suggested that unobserved variables influenced the increase in cell numbers in the untreated group. In contrast, between pre-dose and 24 hr post-dose in the treatment group (400 mg/kg), there was a significant increase in CD4^+^ T cells (*P* < 0.03) and CD8^+^ T cells (*P* < 0.03), but not CD4/CD8 ratio (*P* < 0.80). An example of gating strategy is provided (**Fig S1**). To determine the influence of Alphataxin on cell numbers and exclude the influence of the unobserved variables, the difference in cell numbers between the time points was calculated (post-dose – pre-dose). Comparison of the change in cell numbers (Δ) showed that there was not a significant difference between vehicle control and treatment groups in Δ CD4^+^ (*P* = 0.85) or Δ CD8^+^ T cell numbers (*P* = 0.21), but Alphataxin ftreatment caused a significant increase in Δ CD4/CD8 ratio (*P* < 0.007).

**Figure 1.**
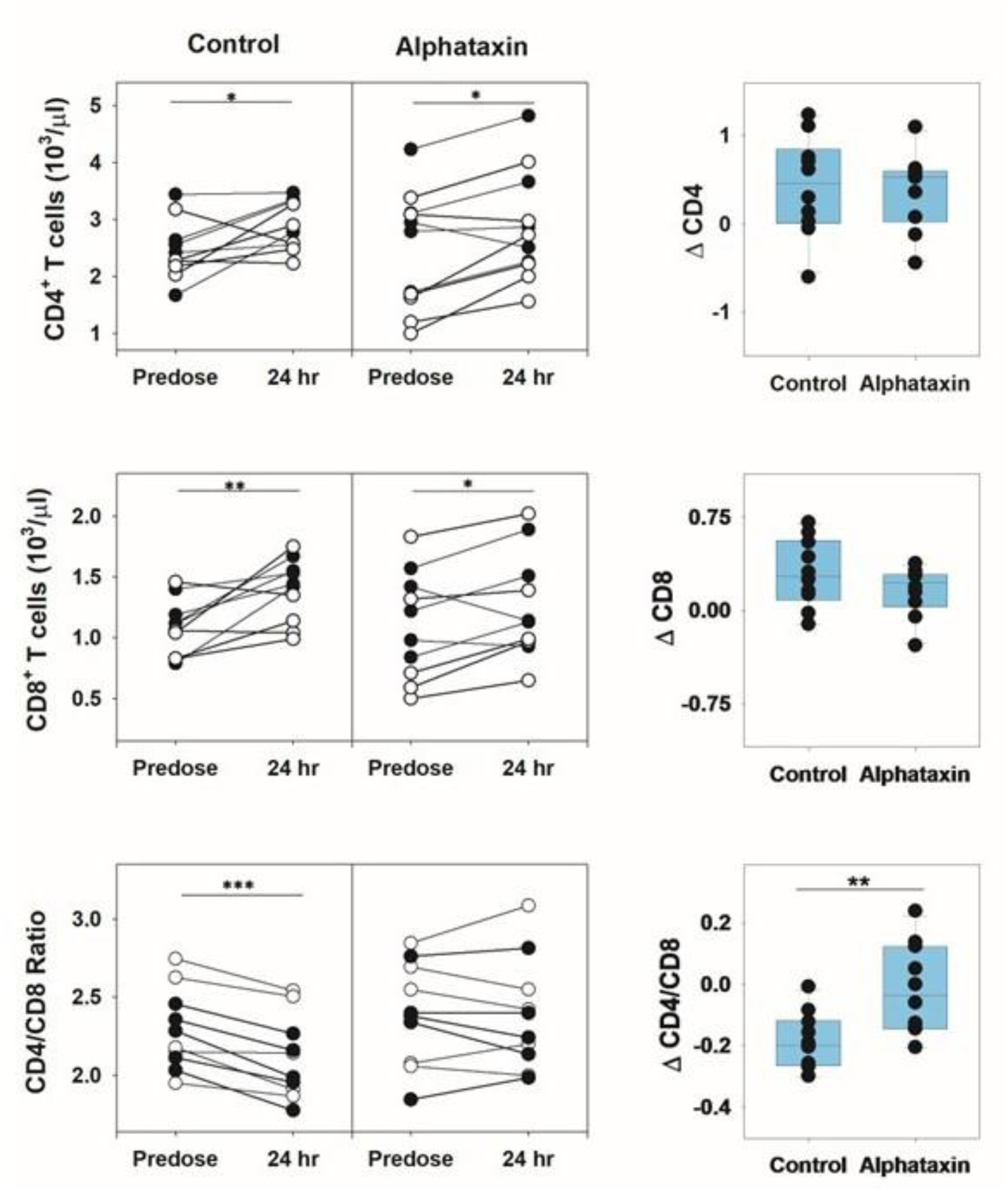
Increased CD4/CD8 ratio 24 hr after single dose of Alphataxin. **A)** Absolute cell counts (cells/µl) were obtained by flow cytometric analysis of blood collected from 5 male (λ) and 5 female (ϒ) rats prior to and 24 hr following oral administration of Alphataxin (400mg/kg). Control rats receiving vehicle (PBS, pH 7.2) by oral gavage exhibited significant changes in mean CD4^+^ T cell numbers (pre-dose = 2.47 ± 0.52, post-dose = 2.89 ± 0.43, *P* < 0.05, *n*=10), CD8^+^ T cells (pre-dose = 1.08 ± 0.23, post-dose = 1.39 ± 0.26, *P* < 0.007, *n*=10), and CD4/CD8 ratio (pre-dose = 2.29 ± 0.26, post-dose = 2.11 ± 0.26, *P* < 0.0002, *n*=10) suggesting the effects of unknown variables. In contrast, rats receiving Alphataxin (400 mg/kg) by oral gavage exhibited significant increases in mean CD4^+^ T cell numbers (pre-dose = 2.58 ± 0.97, post-dose = 2.96 ± 0.96, *P* < 0.03, *n*=10) and CD8^+^ T cells (pre-dose = 1.10 ± 0.44, post-dose = 1.26 ± 0.44, *P* < 0.03, *n*=10), but not CD4/CD8 ratio (pre-dose = 2.40 ± 0.33, post-dose = 2.38 ± 0.35, *P* < 0.80, *n*=10). To remove the effects of the unknown variables, the difference between pre-dose and 24 hr post-dose time points was calculated (Δ cell number = post-dose – pre-dose). Alphataxin treatment exhibited no effect on Δ CD4^+^ T cell numbers (mean = 0.38 ± 0.44, *P* = 0.85, *n*=10) or Δ CD8^+^ T cell numbers (mean = 0.16 ± 0.20, *P* = 0.21, *n*=10) as compared to control rats (mean = 0.43 ± 0.57 and 0.30 ± 0.27, respectively. *n*=10). However, Alphataxin treatment caused a significant increase (*P* < 0.007, *n*=10) in Δ CD4/CD8 ratio (mean = −0.01 ± 0.15) as compared to control rats (mean = −0.18 ± 0.09). An example of gating strategy is provided in Supplementary Material (**Fig S1**).

### Tumor growth is suppressed by Alphataxin treatment in combination with anti-PD-1

Treatment was initiated 3 days after implantation of tumors into the colon (**Fig 2**). The primary comparison was tumor growth in combination-treated animals vs control/monotherapies. Alphataxin was administered daily by oral gavage, and anti-PD-1 was administered twice weekly by IP injection. Tumor size (BLI) was assessed weekly for 31 days (**Fig 3A**). Following 21 days of treatment and 10 additional days of observation following treatment, combination treatment yielded 37.5% ORR. Two clusters of animals were detected and were examined by exploratory bifurcation: those with unabated tumor growth (Combination+, *n*=5), and those with complete or partial response of 97.1% suppression of tumor growth including one animal that achieved tumor-free survival (Combination–, *n*=3) (**Fig 3B**). Tumor size in Combination– treatment was significantly lower than in the Combination+ treatment arm (*P* = 0.04, *n*=8) and the isotype control arm (*P* = 0.02, *n*=7). No significant difference (*P* = 0.60) was detected in BLI between the Combination+ treatment arm and the isotype control arm. The BLI on day 31 suggest that combination treatment, while effective in some mice at the end of the study, may be more effective using higher doses of Alphataxin and anti-PD-1.

**Figure 2.**
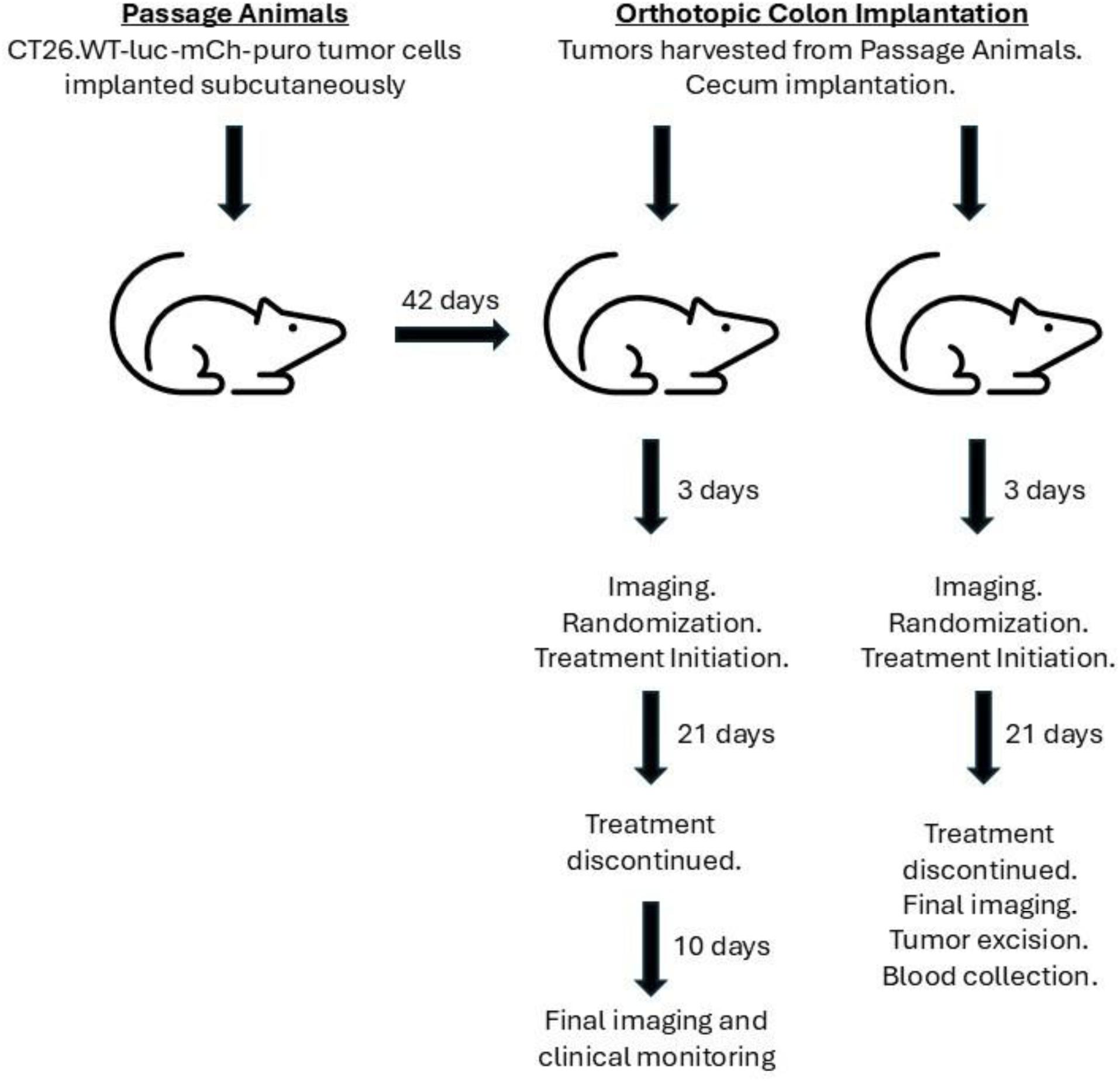
Schematic depicting experimental timelines for Passage Animals and Orthotopic Colon Implantation Animals.

**Figure 3.**
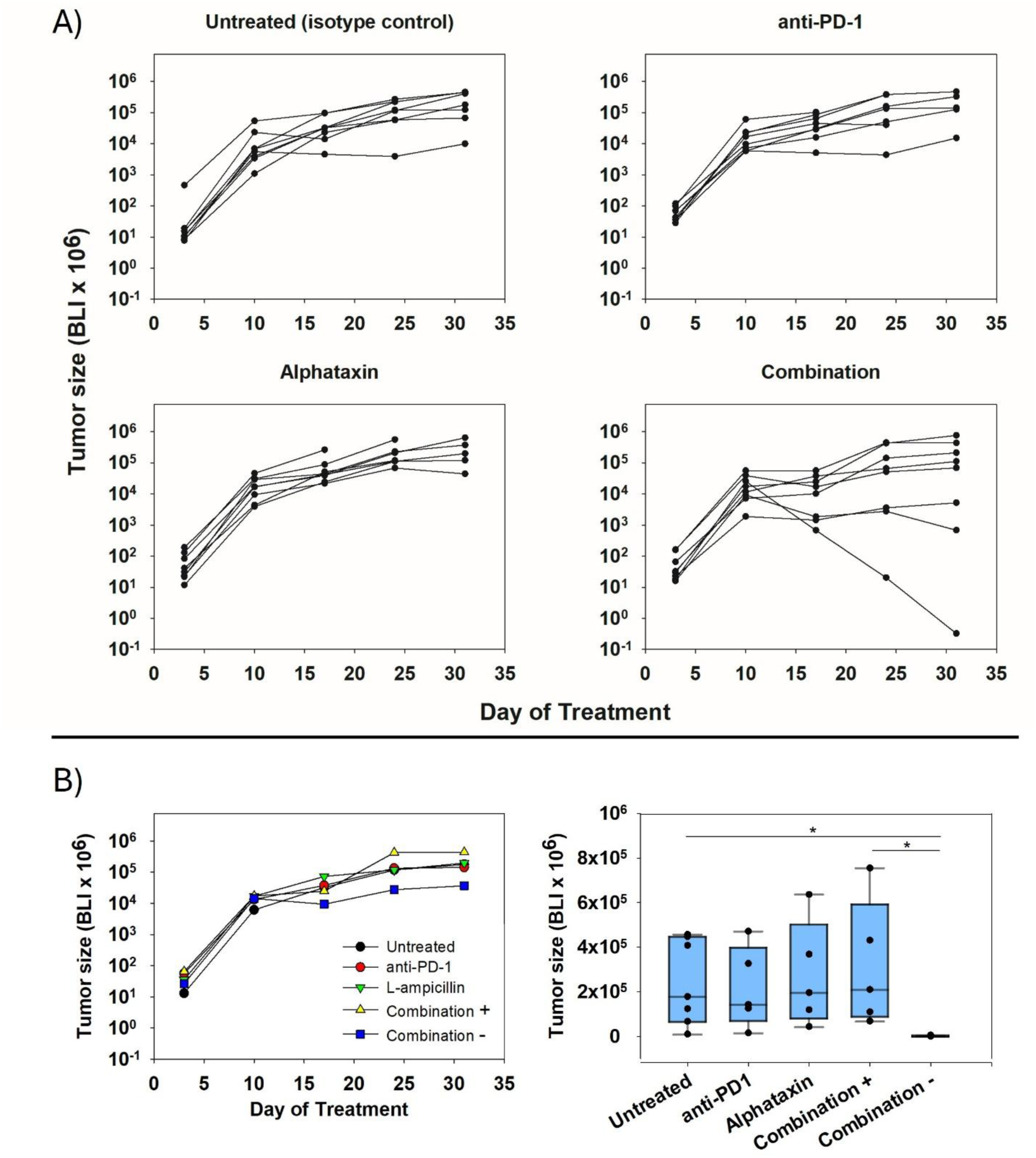
Suppression of tumor growth by combination treatment. Three days after orthotopic tumor implantation, treatment was initiated to include daily oral gavage with Alphataxin and twice weekly IP injection with anti-PD-1 for 21 days followed by 10 days of observation (*n*=8/treatment arm). **A)** Individual tumor growth is depicted through day 31. Surviving mice on day 31 were isotype control (*n*=7), anti-PD-1 (*n*=5), Alphataxin (*n*=5), and combination treatment (*n*=8). **B)** No response was detected in monotherapy arms. The ORR in the combination treatment arm was 37.5%. In mice with tumor growth suppression or regression (Combination–, median BLI = 628 x 10^6^, *n*=3) as compared with isotype control mice (median BLI = 178000 x 10^6^, *n*=7), tumor growth was suppressed 97.1% (*P* = 0.02). In contrast, in combination treatment with no suppression, (Combination+, median BLI = 209,400 x 10^6^, *n*=5), there was no difference as compared with isotype control mice (*P* = 0.60). The mean estimated tumor burden for all groups (*n*=32) in the experiment on the first day of treatment was 6.52E+07 ± 2.2 p/s. Tumor growth kinetics and parameters (clinical signs, body weight change, and time on study) in the untreated arm was within historical norms. ORR = animals responding/total animals treated*100. Asterisks designate statistically significant differences (\**P* < 0.05).

At the last imaging timepoint before any animal exited the study due to disease progression (day 17, *n*=8/arm), animals were evaluated for tumor burden. The median tumor burden on day 17 of Combination– therapy was 4.5% of median isotype control (*P*=0.04) (**Fig 4A**). In contrast, mean tumor burden in Combination+ (76.1%), monotherapy treatments anti-PD-1 (117.2%) and Alphataxin (133.7%) were not significantly different from isotype control (*P* = 0.52). The tumor burden on day 17 suggests that at these doses, monotherapy is less effective than combination therapy in preventing tumor growth.

**Figure 4.**
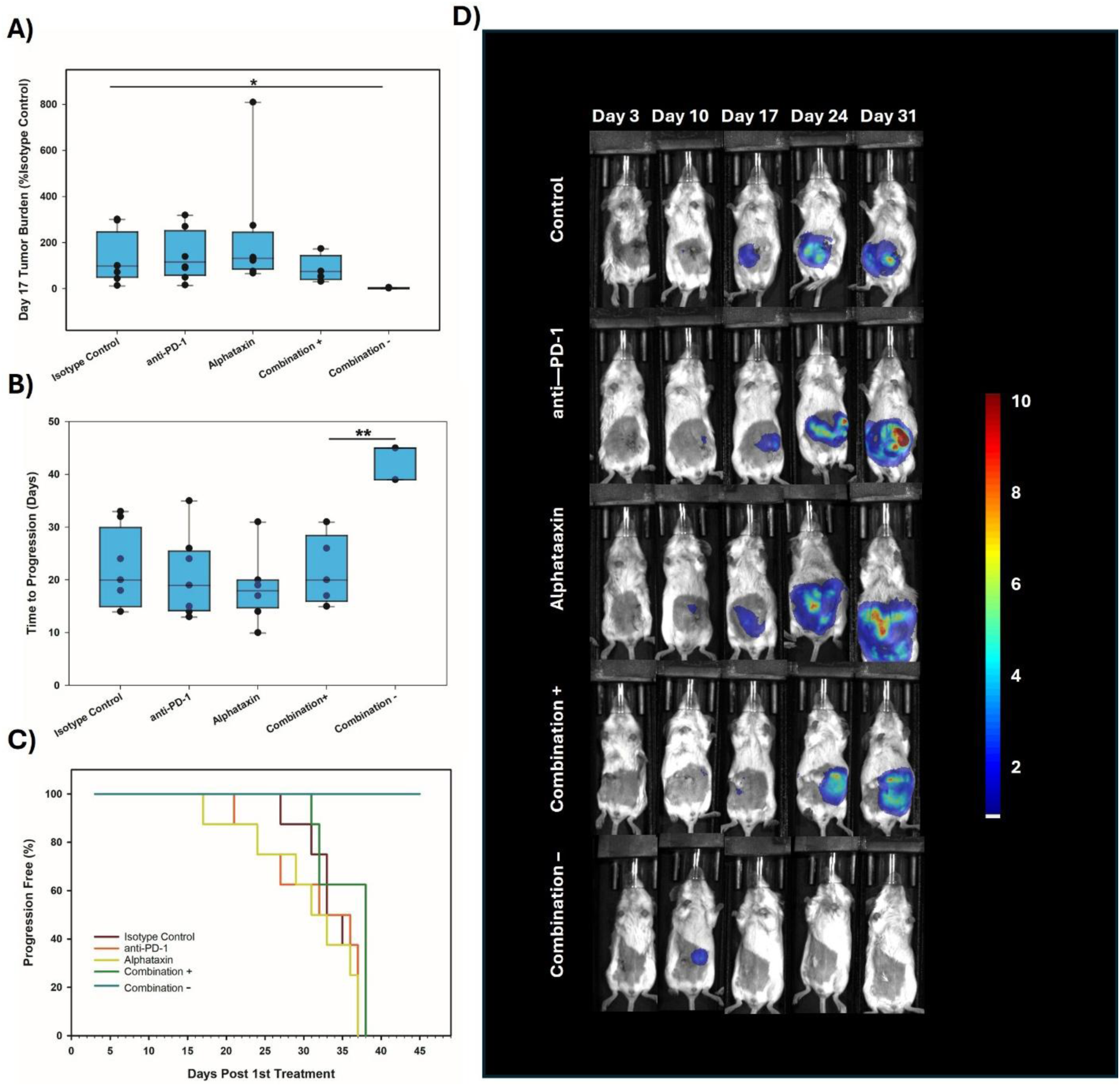
Tumor burden, time to progression, and survival. **(A)** Tumor burden and time to progression were compared on day 17 before any animal exited the study due to disease progression (*n*=8/arm). Combination therapy resulting in suppression or regression of tumor growth (Combination–, *n*=3) resulted in a significantly lower Day 17 median tumor burden of 4.5% of isotype control (*P* = 0.04). Combination therapy with tumor growth (Combination+, *n*=5) resulted in median tumor burden of 76.1%, anti-PD-1 with 117.2%, and Alphataxin with 133.7% of isotype control and were not significantly different (*P* = 0.52, *n*=8/arm). Percent tumor burden is defined as median tumor burden of treatment/ median tumor burden of isotype control * 100 with the median tumor burden of the isotype control arm being 100%. The mean tumor burden for all groups on the first day of treatment was 6.52E+07 ± 2.2 p/s. **(B)** Time to progression in the Combination– arm (median 45 days, *n*=3), including 1 tumor-free survivor, was significantly prolonged as compared with the Combination+ arm (median 20 days, *n*=5, *P* = 0.002). Isotype control (median 20 days, *n*=8), anti-PD-1 (median 19 days, *n*=8), Alphataxin (median 18 days, *n*=8), and Combination+ were not significantly different (*P* = 0.75). (**C**) Survival was significantly extended in Combination– (**turquoise, median survival 45 days**) to the termination of study (*P*<0.001) as compared with isotype control mice (**violet, median 37 days survival**) representing a 22% increase in lifespan [(Treatment – Isotype Control)/Isotype Control*100%]. In contrast, survival in Combination+ (**green, median survival 35 days**), anti-PD-1 (**orange, median survival 35 days**) and Alphataxin (**chartreuse, median 32 days**) arms were not different from Isotype control (*P* = 0.49). **D)** Longitudinal images of the dorsal recumbent supine position of tumor growth after 31 days of treatment. A representative animal was selected that showed similar bioluminescence measurement with at least 2 other animals within the same treatment arm. The mice in the combination treatment arm were bifurcated into 2 groups, those with tumors (Combination+, 4^th^ row, *n*=5) and those without tumors (Combination‒, 5^th^ row, *n*=3). The luminescence color scale represents photons/sec/cm^2^/steradian x 10^9^ with minimum 5e^8^ and maximum 1e^10^. Asterisks designate statistically significant differences (** *P* < 0.01,\**P* < 0.05).

Time to progression as compared with isotype control mice was assessed. In Combination– mice, 12.5% were tumor free (*n*=1) and median time to progression was 45 days in 37.5% of mice (*P* = 0.002, *n*=3). There were no tumor-free survivors in Combination+ with median time to progression of 20 days (*n*=5). Similarly, there were no tumor free survivors in the isotype control arm or monotherapy arms, and median times to progression were 20, 19, and 18 days, respectively, with no significant differences between Combination+, isotype control, or monotherapy (*P* = 0.75) (**Fig 4B**). Time to progression was significantly extended by combination treatment consistent with the ability of combination treatment to prevent tumor growth in some mice at these doses.

As compared with Isotype control mice (median 37 days survival), Combination– mice exhibited significantly extended survival up to the termination of the study at 45 days (*P*<0.001) as compared with isotype control representing a 22% increase in lifespan (**Fig 4C**). Longitudinal images of tumor growth demonstrated that after 31 days of treatment, combination treatment substantially impeded or regressed tumor growth as compared with isotype control or monotherapy treatment (**Fig 4D**). These data demonstrate that combination therapy with Alphataxin and anti-PD-1 can increase survival and potentially allow mice to achieve durable remission. Survival was significantly improved by combination treatment consistent with the ability of combination treatment to increase the time to progression at these doses in some mice.

### Tumor-infiltrating CD4^+^ T cells, CD8^+^ killer T cells, and NK cells are increased by Alphataxin in combination with anti-PD-1

Whereas our previous kidney cancer studies in mice demonstrated that Alphataxin monotherapy significantly increased the intratumoral CD4/CD8 ratio, the present colon cancer study showed no change in CD4/CD8 ratio at termination of the study on day 23 (Bristow *et al*., 2021). However, consistent with Alphataxin’s mechanism of action to mobilize lymphocytes, combination treatment significantly increased (*P*=0.03) the absolute number of intratumoral CD8^+^ T cells (*n*=5) as compared with isotype control (*n*=5) (**Fig 5**). Combination treatment trended toward increased absolute number of intratumoral CD4^+^ T cells (*n*=5) as compared with isotype control (*n*=5) (*P* = 0.1). Combination treatment trended toward increased (*P* = 0.06) intratumoral ratio of CD8/Treg cells (*n*=5) as compared with isotype control (*n*=5), and this effect appeared to be due to Alphataxin treatment (*n*=5) rather than anti-PD-1 treatment (*n*=3). As compared with isotype control (*n*=5), combination treatment significantly increased the absolute number of tumor-infiltrating NK cells (*P* = 0.03, *n*=5), significantly increased the absolute number of M2 macrophages (*P* = 0.009, *n*=5), and trended toward increased absolute number of DC2 dendritic cells (*P* = 0.06, *n*=5). Although there was no difference in the absolute number of CD8^+^ LAG3^+^ cells between combination treatment (*n*=5) and isotype control, there was a significant decrease with anti-PD-1 monotherapy (P = 0.03, *n*=3). These results suggest that the interactions between CD4^+^ and CD8^+^ T cells explain the improved benefit of combination treatment over monotherapies that affect each T cell subtype separately and raise the question whether such interactions might be detected by examining cytokine levels within the tumor.

**Figure 5.**
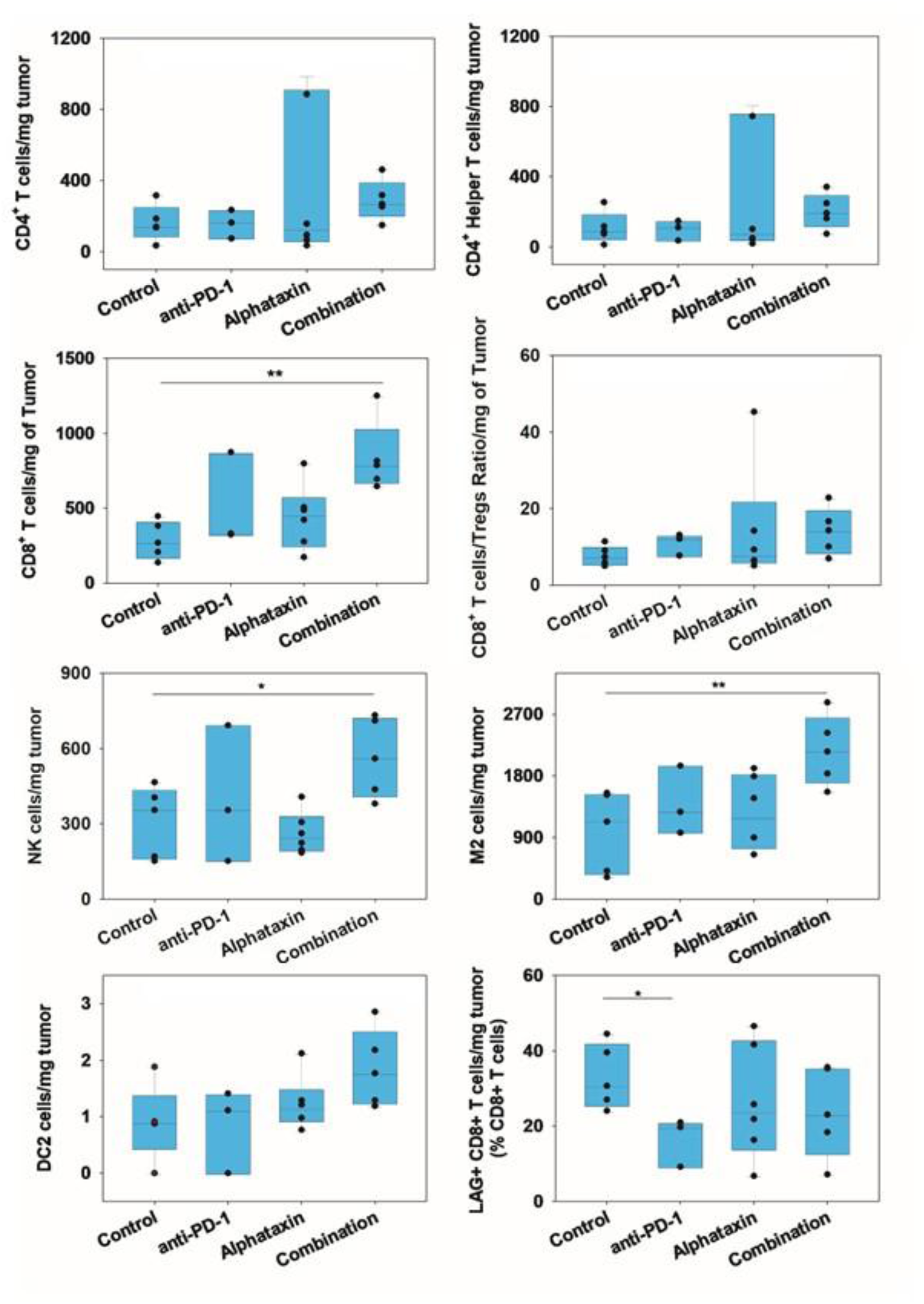
Increased numbers of tumor-infiltrating CD4^+^ T cells, CD8^+^ T cells, NK cells, M2 cells, and DC2 cells. Combination treatment trended (*P* = 0.10) toward increased absolute number of CD4^+^ T cells (median = 289.45 ± 114.06 cells/mg tumor, *n*=5) as compared with isotype control (mean = 161.85 ± 102.38 cells/mg tumor, *n*=5). Combination treatment increased the number of CD4+ helper T cells (mean = 203.88 ± 99.49 cells/mg tumor, *n*=5) as compared with isotype control (mean = 109.99 ± 89.82 cells/mg tumor, *n*=5), but without statistical importance (*P* = 0.15). Combination treatment significantly (*P*=0.03) increased the absolute number of CD8^+^ T cells (median = 786.10 cells/mg tumor, *n*=5) as compared with isotype control (median =268.71 cells/mg tumor, *n*=5). Combination treatment trended (*P* = 0.06) toward increased the tumor-infiltrating ratio of CD8/Treg cells (mean = 14.14 ± 6.12 cells/mg tumor, *n*=5) as compared with isotype control (mean = 7.73 ± 2.55 cells/mg tumor, *n*=5). Combination treatment significantly (*P* = 0.03) increased the absolute number of NK cells (mean = 564.08 ± 157.99 cells/mg tumor, *n*=5) as compared with isotype control (mean = 309.11 ± 141.15 cells/mg tumor, *n*=5), significantly (*P* = 0.009) increased the absolute number of M2 macrophages (median = 2154.75 cells/mg tumor, *n*=5) as compared with isotype control (M2 median = 1129.98 cells/mg tumor), and trended (*P*=0.06) toward increased absolute number of DC2 dendritic cells (mean = 1.86 ± 0.69 cells/mg tumor, *n*=5) as compared with isotype control (DC2 mean = 0.91 ± 0.67 cells/mg tumor, *n*=5). Although there was no difference in the absolute number of CD8^+^ LAG3^+^ cells between combination treatment (mean = 23.88 ± 12.03 cells/mg tumor, *n*=5) and isotype control (mean = 33.15 ± 8.61 cells/mg tumor, *n*=5), there was a significant decrease (*P* = 0.03) with anti-PD-1 monotherapy (mean = 16.61 ± 6.48 cells/mg tumor, *n*=3) as compared with isotype control (*n*=5). Of 6 mice/treatment arm, 1 animal was found dead in the isotype control arm (*n*=5); anti-PD-1 was found to have no tumors in 3 mice (*n*=3); Alphataxin was found to have tumors in all mice (*n*=6); combination treatment was found to have no tumors in 1 mouse (*n*=5). Asterisks designate statistically significant differences (\**P* < 0.05, ** *P* < 0.01). An example of gating strategy from analysis of an animal in the isotype control treatment arm is depicted in Supplementary Material (**Fig S1**).

Cell populations that did not differ in number between treatment groups included Tregs as well as CD8^+^ T cells that were also positive for CD69, PD-1, TIM3, ICOS, Granzyme B, or Ki67. Nor were there differences between treatment groups in the numbers of NK T cells, B cells, CD11^+^ cells, G-MDSC, M-MDSC, M1 tumor-associated macrophages (TAM), DC1 cells, XCR1^+^DC cells, as well as CD4^+^ T cells that were also positive for CD69, PD-1, TIM3, LAG3, or ICOS. In this smaller study, BLI was only measured once per mouse (*n*=6) following treatment initiation precluding comparison of mice with or without tumor suppression.

### IFN-γ^+^ T cells within tumors are decreased by Alphataxin monotherapy

To examine the potential interactions between CD4^+^ and CD8^+^ T cells within the tumors, cytokines associated with specific cell populations were examined. Alphataxin monotherapy significantly decreased detectable IFN-γ (*n*=6) in stimulated tumor-infiltrating CD4^+^ T cells as compared with isotype control mice (*n*=5, *P* = 0.04) (**Fig 6A**). There were no significant changes in IFN-γ in stimulated CD8^+^ T cells as compared with isotype control in any treatment. Examples of gating strategy are depicted (**Fig S2**). These results are consistent with the expected interactions between CD4^+^ and CD8^+^ T cells in the TME.

**Figure 6.**
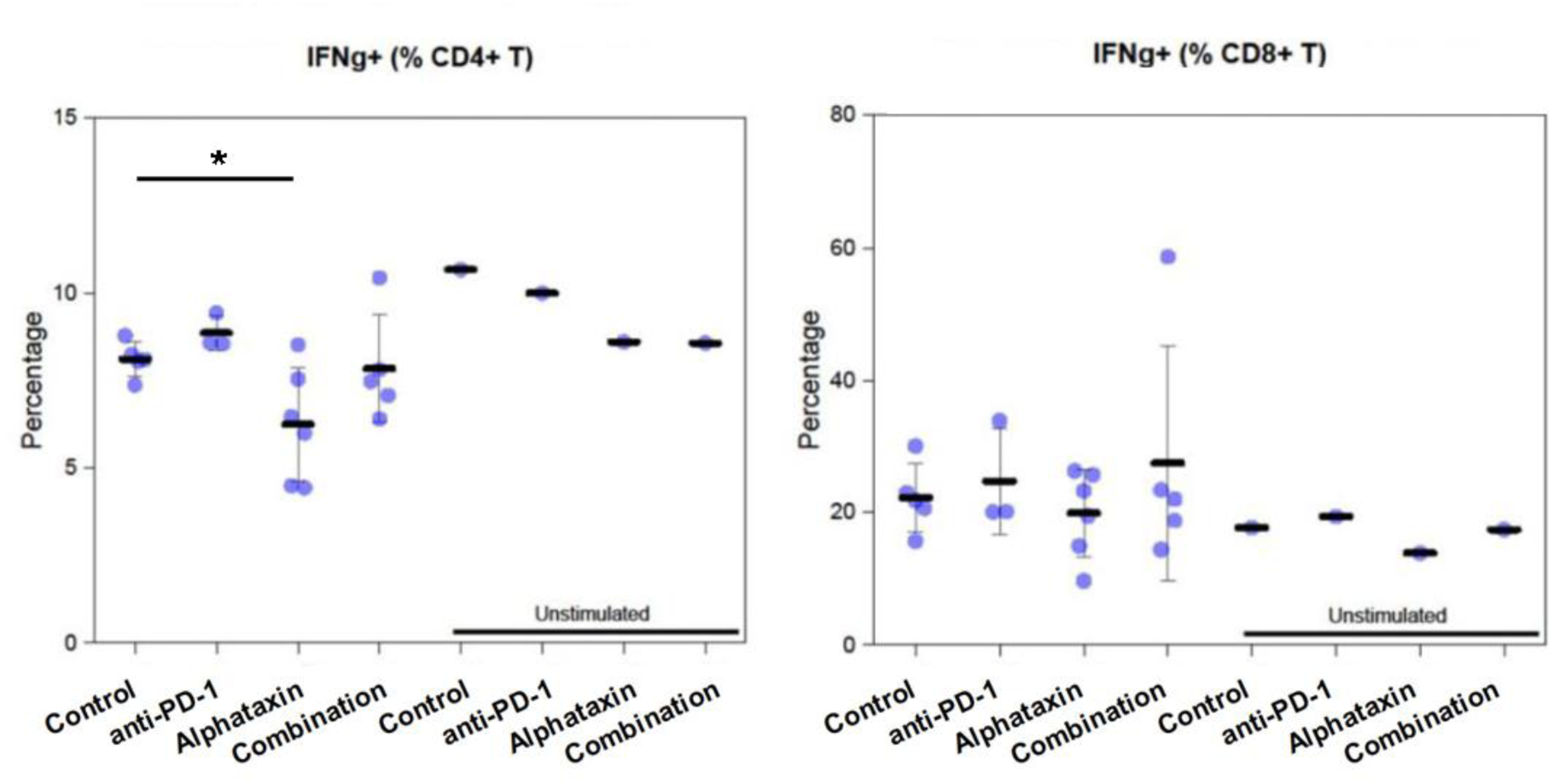
Decreased tumor-associated CD4^+^ T cell IFN-γ. Tumor sampling was conducted on day 23 of treatment as this was the final timepoint for imaging. Cells extracted from tumors were unstimulated or stimulated using PMA/ionomycin/Brefeldin A. **A)** Alphataxin monotherapy significantly decreased detectable IFN-γ (mean = 6.25% ± 1.63% CD4^+^ T cells, *n*=6) in stimulated tumor-infiltrating CD4^+^ T cells as compared with isotype control mice (mean = 8.12% ± 0.50%, *n*=5, *P* = 0.04). In contrast, there was no significant IFN-γ difference between treatments in stimulated CD8^+^ T cells (*P* = 0.87). Of 6 mice/treatment arm, 1 animal was found dead in the isotype control arm (*n*=5); anti-PD-1 was found to have no tumors in 3 mice and insufficient recovery of cells in 1 mouse (*n*=2); Alphataxin was found to have tumors in all mice (*n*=6); combination treatment was found to have no tumors in 1 mouse (*n*=5). Asterisks designate statistically significant differences\**P* < 0.05). Examples of cytokine gating strategy from analysis of tumor cells, tumor cell lymphocytes, and tumor cell monocytic cells and DCs are provided in Supplementary Material (**Fig S2**).

There were no significant changes in TNF-α, IL-2, Granzyme B, and CD107a in stimulated CD4^+^ and CD8^+^ T cells within tumors. The anti-PD-1 monotherapy arm of the tumor cytokine comparison included only 2 of 6 mice as 3 of 6 were reported to have no tumors and insufficient cells were recovered in 1 of the remaining 3 tumor samples. Following treatment initiation, BLI was measured once at study completion precluding comparison of mice with or without tumor suppression.

### Serum inflammatory cytokines are changed by anti-PD-1 treatment

Treatment induced changes in serum cytokine levels were examined for comparison with changes in TME cytokine levels. In contrast to the lack of influence of anti-PD-1 monotherapy on intratumoral T cell cytokine levels, some serum cytokine levels showed changes with anti-PD-1 monotherapy (**Table 1**). For example, in anti-PD-1 monotherapy, as compared with isotype control mice, serum IL-6 was significantly lower (*P* = 0.02), serum IL-1β was significantly lower (*P* = 0.05), serum IL-10 was significantly lower (*P* = 0.04), and serum IL-2 was significantly higher (*P* = 0.04). These results suggest that anti-PD-1 treatment decreased general immune cell activity outside the TME as would be expected.

**Table 1.** Serum inflammatory cytokine levels (pg/ml) in mice treated using Alphataxin and/or anti-PD-1 antibodies. ^1^.

| **Cytokine (pg/ml)** | **Isotype Control (n=5)<sup>2</sup>** | **anti-PD-1 (n=6)** | **Alphataxin (n=6)** | **Combination (n=6)** |
| --- | --- | --- | --- | --- |
| **IFN-γ** | 2.68 ± 1.06 | 5.83 ± 10.29 | 2.19 ± 0.83 | 3.94 ± 2.08 |
| **IL-10** | 34.3 ± 15.6 | **16.3 ± 9.06\*** | 28.3 ± 7.18 | 34.1 ± 16 |
| **IL-12p70** | BLD<sup>3</sup> | BLD | BLD | BLD |
| **IL-1β** | 5.69 ± 1.46 | **3.45 ± 1.7\*** | 5.69 ± 0.95 | 5.12 ± 2.12 |
| **IL-2** | 0.45 ± 0.19 | **1.1 ± 0.55\*** | 0.37 ± 0.16 | **1.13 ± 0.36\*\*** |
| **IL-4** | 0.16 ± 0.11 | 0.52 ± 0.39 | 0.18 ± 0.06 | **0.38 ± 0.06\*** |
| **IL-5** | 2.81 ± 1.94 | 2.88 ± 0.84 | 2.15 ± 1.01 | 3.28 ± 1.37 |
| **IL-6** | 662 ± 625 | **119 ± 140\*** | 509 ± 258 | 1748 ± 2519 |
| **KC/GRO** | 344 ± 205 | 194 ± 209 | 229 ± 104 | 248 ± 142 |
| **TNF-α** | 41.9 ± 23 | 23.1 ± 10.3 | 26.4 ± 3.69 | 29.8 ± 8.57 |

<sup>1</sup> Blood was collected 24 hrs following the final treatment dose (day 23).
<sup>2</sup> One animal found dead on day 22 with autolyzed intestines.
<sup>3</sup> BLD = below limit of detection
<sup>4</sup> Asterisks designate statistical significance (\*\* $P$ < 0.01, \* $P$ < 0.05). Means and standard deviations are depicted.

As compared with isotype control mice, in combination treated mice, serum IL-2 (*P* = 0.004) and serum IL-4 were significantly higher (*P* = 0.01). These results suggest that combination treatment generally activated T cell activity outside the TME.

Unlike anti-PD-1, Alphtataxin monotherapy showed no effects on serum cytokine levels, and this is consistent with its function to induce cellular locomotion without inducting activation.

Serum cytokines IFN-γ, IL-5, KC/GRO, and TNF-α were unchanged by any treatments, and IL-12p70 was below the detection limit in all serum samples. BLI was measured once per mouse (*n*=6) precluding comparison of mice with or without tumor suppression.

### Alphataxin toxicology and pharmacokinetics

No changes were detected in body weight, food consumption, hematology, organ weights, or macroscopic observations following repeat daily dosing of 5 male and 5 female rats for 6 days with the maximum feasible dose of Alphataxin (800mg/kg).

Small molecule Alphataxin was designed for the purpose of binding to cell surface leukocyte elastase (Bristow *et al*., 2021). To examine the behavior of Alphataxin in plasma, liquid chromatography tandem mass spectrophotometry (LCMS/MS) was performed on plasma samples after protein extraction. For standardization, Alphataxin spiked into 100% plasma was detected without any loss of concentration. In contrast, Alphataxin was not detectable in any plasma samples prepared from blood collected at each of various times following *in vivo* oral dosing. The inability to detect Alphataxin in plasma after *in vivo* dosing suggested two possibilities, that Alphataxin was unstable and degraded or that Alphataxin in whole blood was partitioned out of plasma. The *in vivo* biological effects detected here and elsewhere suggested Alphataxin had not been degraded (Bristow *et al*., 2021; Bristow & Winston, 2021a). Since protein had been extracted from plasma prior to LCMS/MS, it was hypothesized that Alphataxin had partitioned with cells *in vivo*. To quantitate how much orally delivered Alphataxin was unstable vs how much had bound to blood cells, Alphataxin was incubated in whole blood containing cells *in vitro* followed by plasma preparation and compared with Alphataxin incubated in plasma lacking the presence of blood cells *in vitro*. At the first timepoint of comparison (time 0), Alphataxin (1.5 mg/ml) spiked into whole blood (**Fig 7A**) was 7.8% the concentration of Alphataxin (1.5 mg/ml) spiked into plasma demonstrating that 92.2% of Alphataxin had rapidly bound to cells (**Fig 7B**).

**Figure 7.**
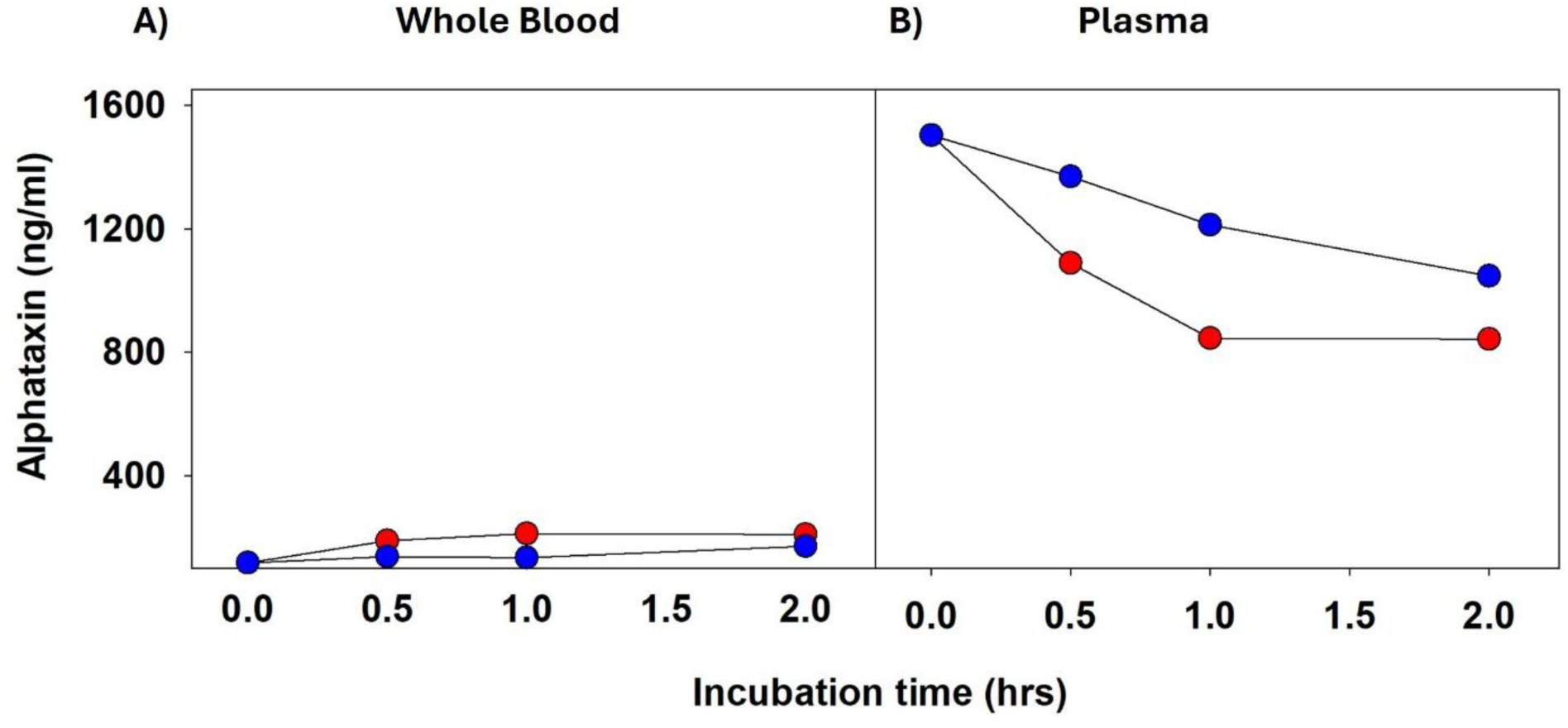
Pharmacokinetics of Alphataxin in whole blood. **A)** Incubation of Alphataxin (1500 ng/ml final concentration) in whole blood at room temperature (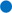) or on wet ice (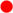) resulted in decreased concentrations detected in subsequently prepared due to its known binding to the cell surface of blood cells. **B)** Incubation of Alphataxin in prepared plasma at room temperature (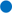) or on wet ice (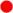) resulted in decreasing concentrations over time due to degradation.

To examine the stability of Alphataxin in plasma and in whole blood *in vitro*, comparisons were made on wet ice and at room temperature. Spiking Alphataxin into whole blood on wet ice, followed by preparing plasma, exhibited a slight increase in detection from 0.117 mg/ml to 0.189 mg/ml after 30 min and 0.212 mg/ml at 60 min that plateaued through 120 min (**Fig 7A**, (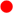). In whole blood, at room temperature, Alphataxin was slightly increased between 0 and 30 min from 0.117 mg/ml to 0.138 mg/ml that plateaued through 60 min followed by a slight increase in detection between 60 min and 120 min (**Fig 7A**, 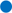). The foremost conclusion from these measurements is that Alphataxin is rapidly bound to cells and is not degraded.

In contrast to spiking whole blood, the concentration of Alphataxin detected after spiking into plasma at room temperature decreased to 1.37 mg/ml (91%) after 30 min and 1.213 mg/ml (80.7%) after 60 min and continued to decrease through 120 min although at a slower rate (**Fig 7B**, 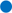). The concentration of Alphataxin detected after spiking directly into plasma on wet ice decreased to 1.089 mg/ml (72%) after 30 min and 0.845 mg/ml (56.2%) after 60 min until reaching a plateau at 60 min through 120 min (**Fig 7B**, 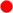). These data suggest that Alphataxin interacts with one or more plasma components differently on wet ice than at room temperature contributing to the differences in stability.

It may be interpreted from these results that Alphataxin is safe, stable for 30 min at room temperature in plasma, and rapidly binds to cell surface leukocyte elastase.

## DISCUSSION

The TME provides a unique platform to investigate the integration of the temporal appearance and disappearance of immune cells and cytokines related to the larger role of the immune system in physiology and pathology as tumors progress or regress in growth. In this study, tumor growth was monitored weekly, but TME was investigated at a single time point, namely at euthanization. Further, the study investigating tumor response (*n*=8) was conducted separately from the study investigating TME (*n*=6) which prevented aligning individual TME with individual growth response. It should be noted that under different treatments, the TME is not strictly comparable due to differences in tumor architecture and growth. Despite these limitations, quantitative comparison of TME cells and cytokines is informative within the context of the tumor progression vs suppression.

The primary findings presented here are that the combination of Alphataxin with anti-PD-1, but not monotherapy at these doses, improves survival, prolongs time to progression, suppresses tumor growth, and in some cases regresses tumors. Importantly, combination treatment was shown to increase the number of tumor-infiltrating CD4^+^ and CD8^+^ T cells as well as NK, M2, and DC2 cells and to decrease the quantity of CD4^+^ T cell-associated IFN-γ. It should be noted that Th1 and Th2 responses differ depending on tumor sites, and this is specifically true for colon cancer which exhibit various CD4^+^ T cell subtypes (Zheng *et al*, 2023). In colon cancer, infiltration with tumor-associated macrophages (TAM) is not associated with poor prognosis whereas in other tumor types, TAM infiltration predicts poor prognosis (Mantovani *et al*, 2022).

IFN-γ is principally released by Th1 CD4^+^ T helper cells, NK, and NKT cells and subsequently binds to receptors present on CD8^+^ killer T cells thereby inducing internalization and stimulation of signaling cascades (Alspach *et al*, 2019). NK and NKT cells are the primary cells in the innate response responsible for releasing IFN-γ and establishing the cytokine milieu that enables the subsequent adaptive immune response in which antigen-specific CD4^+^ Th1 cells release IFN-γ. Diagnostic IFN-γ release assays (IGRA) were developed as an aid for detecting infection, particularly *Mycobacterium tuberculosis*. IGRA are especially useful in immunocompromised individuals, such as those with advanced HIV infection (Mazurek *et al*, 2010). Exposure of T cells to certain antigens *in vitro* produces increased IGRA, and the increased release of IFN-γ in the assay is interpreted to originate from Th1 helper T cells. In contrast, a decrease in detectable IFN-γ and TNFα *in vivo* is interpreted to be due to the uptake of IFN-γ and TNFα by CD8^+^ killer T cells. For example, it has been demonstrated that in COVID-19 positive individuals (*n*=75), IGRA values were significantly lower than in COVID-19 negative individuals (*n*=105) (Coppock *et al*, 2022). This suggests that in COVID-19 positive individuals, IFN-γ released by CD4^+^ T cells *in vivo* may have been bound and internalized by functional immune cells, notably CD8^+^ T cells (Rha & Shin, 2021). Similarly, in the present study, decreased IFN-γ detected in the tumor-infiltrating CD4^+^ helper T cells after Alphataxin monotherapy suggests that the tumor-infiltrating Th1 helper T cells had released IFN-γ. In contrast, IFN-γ levels in stimulated cells from isotype controls, anti-PD-1, or combination treated arms were not different from pooled unstimulated intratumoral CD4^+^ T cells.

IFN-γ has multiple effects within tumors where it activates antigen-specific CD8^+^ killer T cells, establishes a positive feedback loop for its own production by antigen-specific CD4^+^ Th1 cells, inhibits Th1 cells from differentiation into other subclasses, skews differentiation toward Th1 phenotype, and inhibits protumorigenic effects of regulatory T cell (Treg) activity, all of which facilitate suppression of tumor growth (Alspach *et al*., 2019; Lee *et al*., 2021). Here we found that the CD8/Treg ratio was increased by combination treatment consistent with expected IFN-γ effects. It can be interpreted from these results that the tumor-infiltrating CD4^+^ T cells that were increased by Alphataxin treatment were antigen- specific and that the change in IFN-γ levels in tumor-associated CD4^+^ helper T cells contributed to tumor suppression.

In contrast to activation of CD8^+^ killer T cells and tumor suppression, prolonged IFN-γ exposure acting on macrophages and dendritic cells (DCs) may promote tumor cell growth, although DCs are a small fraction of tumor-associated immune cells (Gocher *et al*, 2022; Ivashkiv, 2018; Mellman *et al*, 2023).

The discrepancies between different tumor types regarding immune cell subtype infiltration and cytokine release provide confounding TME scenarios for interpreting the effects of therapeutics. The TME evolves as the tumor and immune response evolves complicating interpretations of the fluidity of tumor growth and suppression, the influence of lymphocyte and other immune cell phenotypes, and the influence of cytokines (Giles *et al*, 2023). This fluidity, or “cancer-immunity cycle,” involves the full physiology of the cancer patient including the patient’s microbiome. The intriguing results discussed herein call for kinetically measuring, in addition to endpoint measuring, of tumor infiltrating cells and cytokines in individual animals to better understand the integrating and modulating roles of specific factors that promote tumor regression.

While Alphataxin has been previously shown to elevate CD4/CD8 ratios following 21 days of treatment, the rapid elevation of CD4/CD8 ratios after a single dose suggests that Alphataxin mobilizes mature T cells from tissue in addition to mobilizing immature T cells within thymus (Bristow *et al*., 2021). Despite the fluidity of the immune response within tumors, it has now been shown that Alphataxin, at its minimum effective dose (10mg/kg) enhanced the efficacy of anti-PD-1 suggesting that Alphataxin at a dose level closer to its maximum feasible dose of 800mg/kg in a greater number of animals may produce a more favorable ORR providing more robust augmentation of anti-PD1 efficacy and, more importantly, to expand the number of patients who respond to anti-PD-1 therapy.

## CONCLUSIONS

The alpha-1 antitrypsin surrogate, orally-available Alphataxin, to our knowledge, the first and only drug developed to rapidly and sustainably increase the number of circulating and tumor-infiltrating CD4^+^ helper T cells, is a powerful therapeutic that provides long-term remission in T cell-responsive cancers in mice in combination with anti-PD-1.

## METHODS

### Reagents

Alphataxin (CAS# 19379-33-0) was chemically synthesized (Richman Chemicals, Spring House, PA, USA) as previously described (Rolinson *et al*, 1960). A solution of Alphataxin in Dulbecco’s phosphate buffered saline (DPBS) was administered to mice daily for 21 days by oral gavage (10mg/kg in 0.25ml). Monoclonal anti-PD-1 antibody (Clone RMP1-14) or isotype control (Clone 2A3) (BioXCell, West Lebanon, NH, BE0146) were administered to mice twice weekly for 21 days via intraperitoneal (IP) injection (10mg/kg in 0.01 0.25ml DPBS).

### Animal studies

Animal studies were performed by LabCorp Early Development Laboratories Inc. under approved IACUC VUF #73: Abdominal Orthotopic Tumor Models for Evaluation of Cancer Therapies. Female Balb/c mice were obtained from Envigo at 5-6 weeks of age and housed in Labcorp’s facilities. Toxicology and pharmacokinetics studies were performed under IACUC-approved protocols 8498821 and 8498822. Male and female Sprague Dawley rats were obtained from Charles River Laboratories at 6-8 weeks of age. Animals were euthanized by determination of the Labcorp Development’s veterinarian using CO_2_ exposure with flow rate to provide 30-70% displacement if demonstrating weight loss >20%, severe diarrhea, severe and continued signs of pain (rough hair coat, hunched/tucked posture, writhing, loss of nesting interest, lethargy), ileus abdominal distension, severe/progressive infection or wound closure failure at the surgical site, severe respiratory distress, loss of righting reflex, or tumor burden >10% of animal’s body weight. It was not appropriate or possible to involve patients or the public in the design, or conduct, or reporting, or dissemination plans of our research

### Pharmacokinetics and toxicology

#### Dose Justification

In a previous dose escalating study in mice, the dose levels of 100, 200, 300, and 400 mg/kg were well tolerated without any clinical findings or changes in body weight or food consumption. Thus, the maximum tolerated dose was not established in the previous mouse study, and the starting dose for Phase I of this rat study was selected at 100 mg/kg. Phase II (DRF) of the study was determined based on findings in Phase I.

#### Justification for Number of Animals

For Phase I (Single Dose Phase), the number of animals (3/sex/group) is considered to be the appropriate and minimum number necessary for generating relevant data to evaluate potential test article-associated toxicity and expected variability among animals. Results of this phase were used to support dose selection for the subsequent repeat dose phase. For Phase II (Repeat Dose Phase), the number of animals in the protocol is considered to be the minimum necessary for statistical, regulatory and scientific reasons.

The number of animals selected for toxicokinetic evaluations (6/sex/treated group) is considered the minimum number necessary to provide meaningful data, given the inherent variability in absorption, distribution, metabolism and excretion processes. A control group with 3 animals/sex is necessary to evaluate the absence of the test article. Multiples of the number of animals for each time point were specified based on the volume of blood required for the toxicokinetic assay and the limited volume of blood which can be obtained from a rat.

Medical treatment necessary to minimize suffering and distress, including euthanasia, was the sole responsibility of the attending laboratory animal Veterinarian.

Because Alphataxin is known to bind to the cell surface, pharmacokinetics was performed either by spiking Alphataxin into plasma or by spiking into whole blood (Bristow & Winston, 2021a). Whole blood was separated into plasma or cells by centrifugation. For spiking, plasma or whole blood were incubated with Alphataxin on ice or at 23°C for 0, 30, 60, or 120 minutes. Samples were maintained at −20°C until quantitation using liquid chromatography tandem mass spectrophotometry (LCMS/MS).

### Liquid chromatography

Liquid chromatography to detect Alphataxin in plasma was performed using a Phenomenex Synergi Polar-RP 80Å column (50× 3 mm, 4 µm particle size). Methanol:water:ammonium hydroxide (50:50:0.1) was prepared daily using HPLC grade methanol, Super-Q, type 1 HPLC grade water, and ACS reagent-grade ammonia hydroxide. Methanol:water:formic acid (50:50:0.1) was prepared daily using ACS reagent-grade formic acid. Acentronile: Methanol (50:50) was prepared in advance for protein extraction.

Alphataxin (approximately 2.75 mg) was dissolved in approximately 2 ml freshly prepared Methanol:Water:Ammonium hydroxide (50:50:0.1), sonicated until completely dissolved (approximately 5 min), and immediately neutralized by adding approximately 0.750 ml freshly prepared methanol/water/formic acid (50:50:0.1) to achieve a concentration of 1 mg/ml. Single-use aliquots of 250 µl were stored at −70°C. Internal standard, Alphataxin-d5, was prepared similarly.

Calibration standards were prepared using Alphataxin at concentrations of 50 to 50,000 ng/ml. QC samples were prepared from separate stock solutions of Alphataxin.

Samples (n=3 per time point) were prepared by aliquoting 25 µl per well into a 96 well plate. To the blank (water) was added 25 µl methanol:water (50:50). To all other wells, was added 25 µl internal standard solution. To all wells was added 300 µl Acetonitrile:Methanol (50:50). After sealing and vortexing (3 min) the plate was centrifuged at 1640 x g for 5 min. at 4°C. To a clean 96 well plate, 50 µl supernatant was transferred, and 300 µl water was added to each well. The plate was sealed and vortexed for 3 min. prior to analysis.

Analysis was conducted using column temperature of 40°C and autosampler temperature of 5°C. Mobile Phase A was water:formic acid (100:0.1) and Mobile Phase B was methanol:formic acid (100:0.1). Injection volume was 1-5 µl with backpressure typically 97 bar and total flow of 0.7 ml/min.

### Orthotopic implantation of CT26 colon tumor cells

#### Passage mice

Mice were implanted subcutaneously in the high axilla with CT26.WT-luc-mCh-Puro cells (10^6^ cells/implant in 200µL). CT26-luc-mCh-Puro tumor fragments (approximately 2mm^3^) were obtained from the subcutaneous tumors of the passage animals when a tumor volume reached approximately 500mm^3^ (42 days). Necrotic areas were discarded.

#### Orthotopic implant mice

The implant site was shaved 48 hours prior to surgery. An incision was made in the peritoneum so that the cecum could be gently pulled through and exposed. A tumor fragment (2mm^3^) was adhered to the cecum using sutures in a figure eight pattern. The peritoneum was closed using absorbable sutures. The skin was closed using absorbable sutures and stainless steel wound clips.

### Bioluminescence (BLI) imaging of tumor size

Scanning for bioluminescence intensity (BLI) was performed using IVIS Spectrum (Perkin Elmer) 3 days following implantation on animals in the dorsal recumbent supine position. Animals were distributed into treatment groups based on tumor burden calculated from BLI data such that the mean BLI in each group was within 10% of the overall mean (*n*=8 mice/group for efficacy and *n*=6 mice/group for tumor immunophenotyping and cytokine analysis). BLI background signal for this study was measured at 6.20E+05 p/s.

For comparison of treatment efficacy, images were acquired weekly on days 10, 17, 24, 31, 38, 45 to obtain whole body bioluminescence. Animals were held for survival endpoints with a maximum of 45 days, and tumor growth suppression was determined based on BLI. Efficacy was expressed as objective response rate (ORR) calculated as number of animals exhibiting complete or partial tumor growth suppression divided by number of animals treated. For TME analysis, images were acquired on days 3 and 23 to obtain whole body bioluminescence.

### Serum cytokine analysis

Mice were euthanized by CO_2_ exposure 24 hr following the final treatment dose (day 23), and full blood volume was collected by cardiac puncture. Serum was processed for cytokine analysis (IFN-γ, IL-1β, IL-1, IL-4, IL-5, IL-6, IL-10, IL-12p70, KC/Gro, and TNF-α) using the V-PLEX Pro-inflammatory Panel 1 Mouse kit (Meso Scale Diagnostics, Rockville, MD) using Quickplex SQ 120 equipment. Detected cytokines are expressed as the quantity within the gated cells (% cells).

### Flow cytometry for tumor-infiltrating cells and associated cytokines

Immediately after blood collection, tumors were excised and processed into single cell suspensions for flow cytometric analysis using the Expanded CompLeukocyte Panel 1 (CD45, CD3, CD4, CD8, FoxP3, CD25, Ki-67, CD69, PD-1, LAG-3, TIM-3, ICOS, CD49b, CD335, Granzyme B), Expanded CompLeukocyte Panel 2 (CD45, CD11b, Ly-6G, Ly-6C, CD11c, CD24, F4/80, MHC Class II, CD206, CD103, XCR1, CD19), and the Cytokine/Granzyme B panel (Labcorp Early Development Laboratories Inc.). Antibodies used for cell phenotype detection were proprietary. The Cytokine/Granzyme B panel (CD45, CD3, CD4, CD8, CD49b/CD335, IFN-γ, TNF-α, IL-2, Granzyme B, and CD107a) was used to compare unstimulated tumor cells with tumor cells stimulated *in vitro* with PMA/ionomycin and Brefeldin A. Detected cells were expressed as the number of cells per mg tumor tissue sampled. Examples of gating strategy are provided in Supplementary Material.

Quality check of the flow cytometry data revealed that there were sufficient events for analysis in all samples acquired. Flow cytometric analysis was judged to be satisfactory based on Fluorescence Minus One (FMO) controls for all fluorochromes used in the setting of gates. Samples with fewer than 10,000 live cells as determined by the viability gate were excluded from analyses. Cells counts represent a relative measure of events collected per sample (typically 10^6^ cells). Absolute counts were calculated as percentage of the number of CD45^+^ cells in each cell suspension by counting using Precision Count Beads (BioLegend) yielding the absolute number of each cell type per gram of tissue.

### Statistical analysis

Means were compared using one-way analysis of variance or Student’s t-test when data were normally distributed. Medians were compared using one-way Kruskal-Wallis analysis of variance on ranks or the Mann-Whitney rank-sum test when data were not normally distributed. Paired t-test was used for pre-and post-dose comparisons. Statistical comparisons and graphs were performed using SigmaPlot software with a significance level of alpha = 0.05 and sufficient power of test (0.8 unless stated otherwise) for all reported comparisons. Error bars represent standard deviations in all cases.

## DATA AVAILABILITY

The data generated in this study are available upon request from the corresponding author.

## AUTHOR CONTRIBUTIONS

**CLB:** Conceptualization, data curation, formal analysis, funding acquisition, investigation, methodology, project administration, resources, supervision, validation, writing-original draft, writing-review and editing. **TQG:** Conceptualization, informal analysis. **RW:** Funding acquisition, resources, supervision.

## ACKNOWLEDGEMENTS

The authors gratefully acknowledge the efforts and advice from Labcorp Early Development Laboratories Inc. and structure analysis by the Dept. of Chemistry, Stony Brook Univ. Funding was generously provided to CLB by the Harry Winston Research Foundation which had no role in the study design, collection, analysis, and interpretation of data, writing of the report, or decision to submit the paper for publication.

